# Derivation of a simple postoperative delirium incidence and severity prediction model

**DOI:** 10.1101/426148

**Authors:** Lindroth H., Bratzke L., Twadell S., Rowley P., Kildow J., Danner M., Turner L., Hernandez B., Chang W., Brown R., Sanders R.D.

## Abstract

**Background:** Delirium is an important postoperative complication, yet a simple and effective delirium prediction model remains elusive. We hypothesized that the combination of the National Surgical Quality Improvement Program (NSQIP) risk calculator for serious complications (NSQIP-SC) or risk of death (NSQIP-D), and cognitive tests of executive function (Trail Making Test A and B [TMTA, TMTB]), could provide a parsimonious model to predict postoperative delirium incidence or severity.

**Methods:** Data were collected from 100 adults (≥65yo) undergoing major non-cardiac surgery. In addition to NSQIP-SC, NSQIP-D, TMTA and TMTB, we collected participant age, sex, ASA score, tobacco use, type of surgery, depression, Framingham risk score, and preoperative blood pressure. Delirium was diagnosed with the Confusion Assessment Method (CAM), and the Delirium Rating Scale-R-98 (DRS) was used to assess symptom severity. LASSO and Best Subsets logistic and linear regression were employed in line with TRIPOD guidelines.

**Results:** Three participants were excluded due to intraoperative deaths (2) and alcohol withdrawal (1). Ninety-seven participants with a mean age of 71.68±4.55, 55% male (31/97 CAM+, 32%) and a mean Peak DRS of 21.5±6.40 were analyzed. Of the variables included, only NSQIP-SC and TMTB were identified to be predictors of postoperative delirium incidence (p<0.001, AUROC 0.81, 95% CI: 0.72, 0.90) and severity (p<0.001, Adj. R^2^: 0.30).

**Conclusions:** In this cohort, preoperative NSQIP-SC and TMTB were identified as predictors of postoperative delirium incidence and severity. Future studies should verify whether this two-factor model could be used for accurate delirium prediction.

## Introduction

Delirium, an acute brain failure, is a common surgical complication experienced by approximately 50% of patients, incurring an estimated annual U.S. cost of $152 billion.^1-4^ As such, delirium is a crucial public health concern, as it is significantly associated with increased mortality^5^ and morbidity^6^, subsequent cognitive decline,^7, 8^ and loss of independence.^5, 9^ One in three cases of delirium may be preventable when multicomponent delirium prevention measures are implemented.^10, 11^ Implementation of such measures would likely be most efficient in high risk individuals, yet clinicians are encumbered by an arduous list of potential patient and perioperative risk factors to identify at risk individuals. Prediction models facilitate the identification of high-risk individuals with the area under the receiver operator curve (AUROC) statistic, which is used to evaluate predictive ability to differentiate health and disease, with a value of 0.5 being no better than chance and 1.0 being perfect. Our recent systematic review identified moderate predictive ability (AUROC 0.73-0.94) in four postoperative delirium prediction models.^12^ While an AUROC of 0.94 is excellent, that model includes postoperative data, which may be inflating model performance, thus invalidating its use as a *preoperative* prediction model. Critically, we did not identify a prediction model for delirium severity in the literature. A severity model may have the most utility, as it would identify patients who are likely to incur the severest delirium and therefore are most likely to benefit from a delirium prevention plan. An ideal model that is clinically applicable would be predictive of delirium incidence and its severity as well as parsimonious, such that it would use a limited number of variables that can be cheaply and easily obtained.

The ability to identify high-risk individuals prior to their surgery is critical to delirium prevention. To identify potential candidate predictors, we considered the pathogenesis of delirium and sought to identify both predisposing (age, depression, and medical comorbidities) and precipitating (surgery) factors. The *Cognitive Disintegration Model* posits that an individual with increasing risk, or vulnerability, to delirium will require less of a precipitating stimulus to cross over the “Delirium Threshold” and become delirious.^13^ This is illustrated in the cognitive trajectory in Figure 1. In contrast, an individual with less predisposing risk factors will require a large stimulus to precipitate delirium. Hence, it is crucial to consider the future precipitating event into delirium prediction models when possible. The National Surgical Quality Improvement Program (NSQIP) online risk calculator^14, 15^ combines several comorbidities (that provide information on predisposing risk factors) and the estimated magnitude of the precipitating event, the surgery, to calculate risk of events. The NSQIP risk score has been widely validated and applied to predict outcomes in various surgical populations, but has not been applied in delirium prediction.^12, 16-19^ We hypothesized that NSQIP risk of serious complications (NSQIP-SC) would be a stronger predictor of delirium over NSQIP risk of death (NSQIP-D) as the causal relationship between serious complications and delirium is likely stronger than the association between delirium and the risk of death.^20^ As delirium is a cognitive disorder, we further hypothesized that cognitive data (that is not included in NSQIP scores) could enhance the prediction of the surgical risk scores. Indeed our recent systematic review identified that current delirium prediction models do not evaluate specific cognitive domains, such as executive function.^12^ Executive function facilitates attention and problem-solving.^21^ Significant associations between preoperative executive function and postoperative delirium incidence have been reported.^22-24^ In sum, our aim was to examine the predictive ability of the NSQIP risk scores with a measure of executive function among other potential delirium risk factors to develop a parsimonious model to predict postoperative delirium incidence and severity.

**Figure 1.**
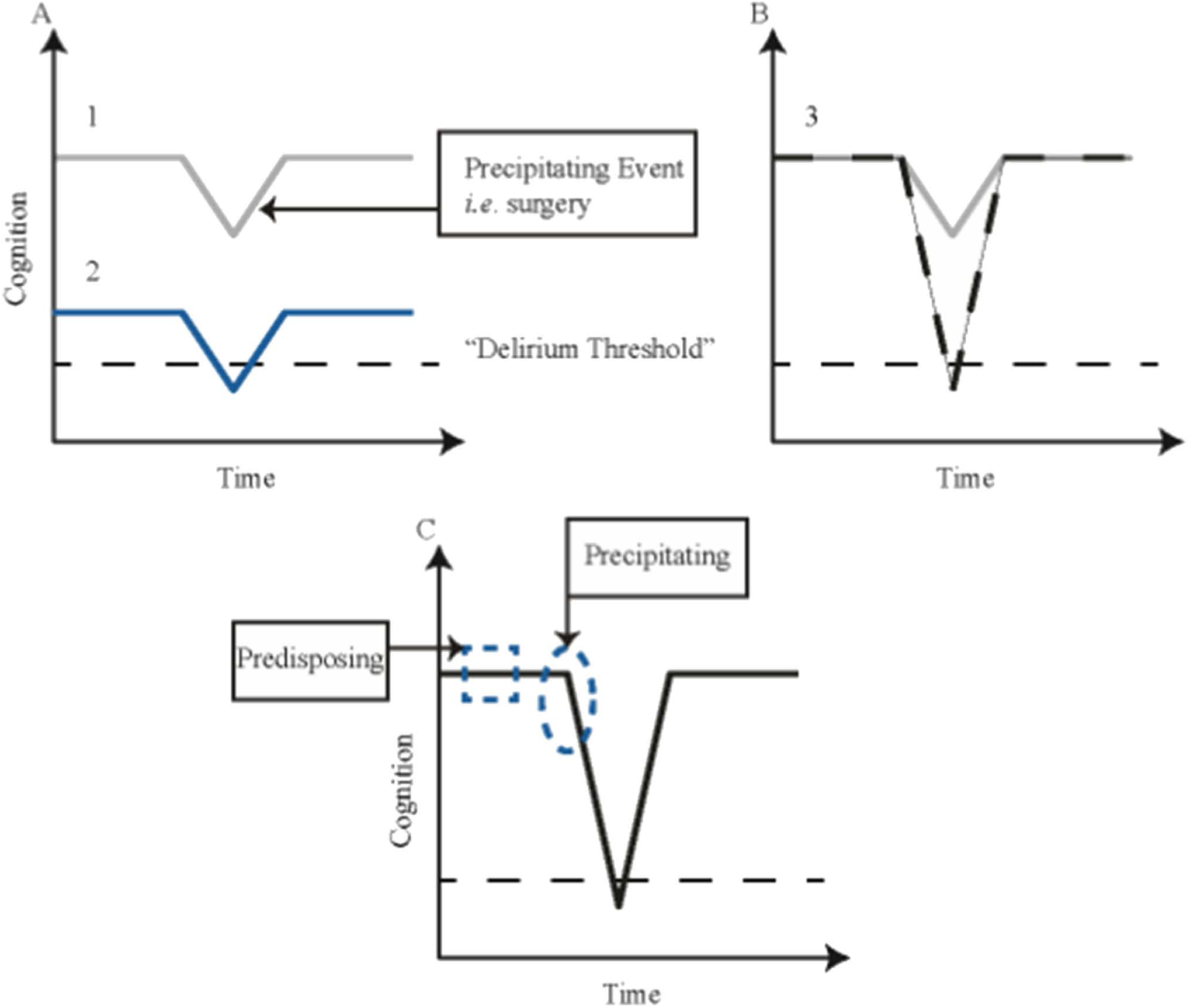
Illustrates the Cognitive Trajectory. The relationship between cognitive abilities (predisposing, y axis) and the precipitating event, *i.e.* surgery over time (x-axis) is shown, with each individual trajectory displayed with a horizontal line. The dashed line, situated above the x-axis of “time”, represents the “Delirium Threshold.” (A) Trajectory #1 (gray line, numbered 1) displays an individual with maximum cognitive abilities. They have a surgery, but do not cross the “Delirium Threshold.” Trajectory #2 (blue line, numbered 2) contrasts #1 by showing an individual with decreased cognitive abilities. This individual undergoes the same surgery and crosses over the “Delirium Threshold” to experience delirium. (B) Trajectory #3 (black-dashed line, numbered 3) returns to an individual with maximum cognitive abilities. A sufficiently large precipitating event will push this individual across the “Delirium Threshold”, inducing delirium. Trajectory #1 (gray line) is transposed onto this graph to show the difference in magnitude and impact of the precipitating event. (C) When developing a prediction model for delirium, it may be important to consider not only the predisposing risk factors, but also the influence of the precipitating event. A surgical risk score such as NSQIP combines both predisposing risk and the future-precipitating event into one score, which may be optimal for postoperative delirium prediction.

## Methods

This analysis is a sub-study drawn from an ongoing prospective perioperative cohort study that is approved by the University of Wisconsin-Madison, Health Sciences Institutional Review Board (#2015-0374) and registered with ClinicalTrials.gov (ref: NCT03124303, NCT01980511). Between August 2015 and May 2018, 1,049 potential participants were screened from vascular, urology, general and spine surgical clinics (HL, ST, LT, PR, JK, MD, BH, RDS). As shown in Figure 2, 100 subjects were recruited and 97 were included in the final analysis.

**Figure 2.**
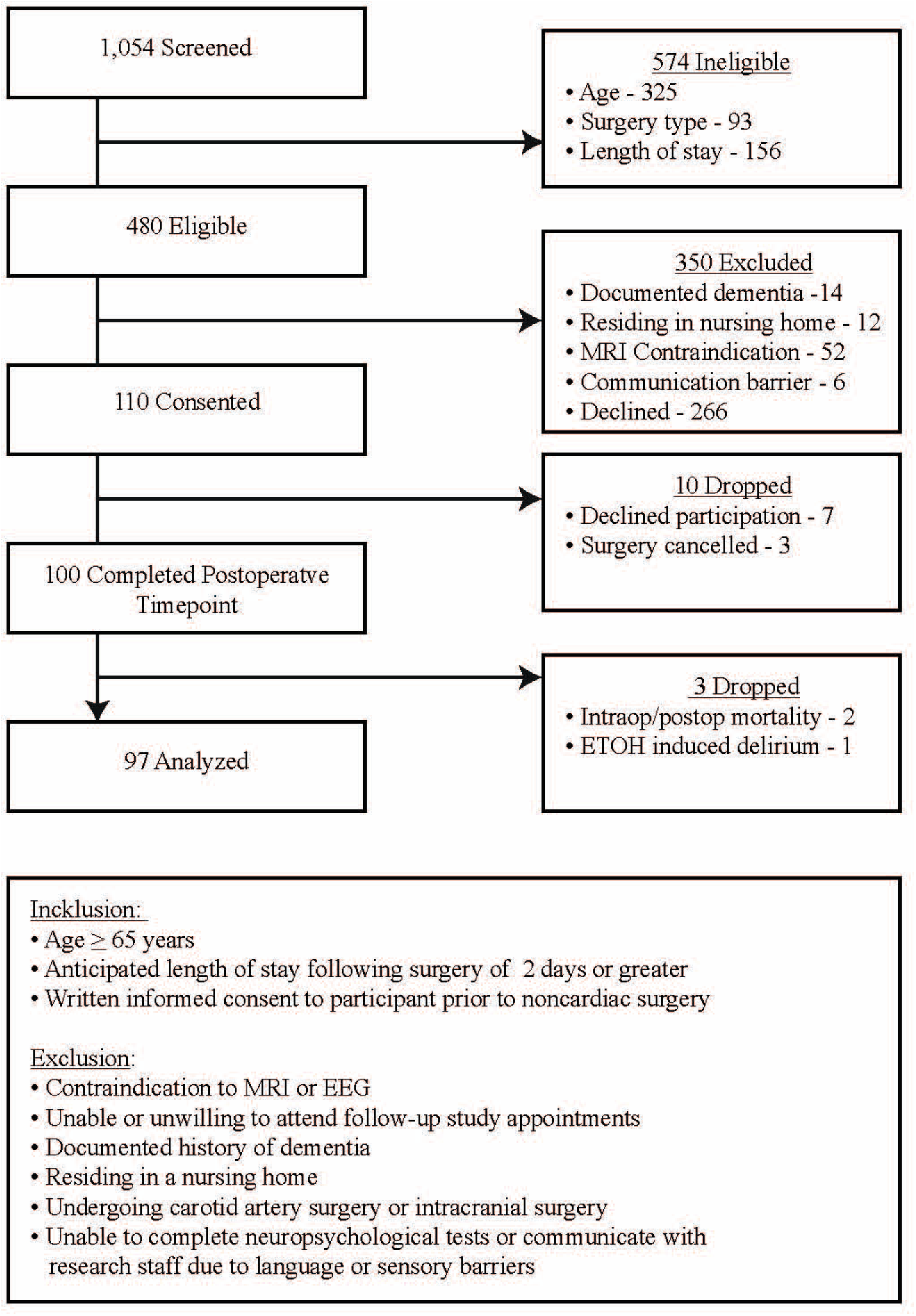
Displays the inclusion and exclusion criteria and a flowchart detailing study screening, recruitment, consent, and attrition numbers.

### Preoperative Predictors and Assessment

Preoperatively, participants underwent an interview and completed assessments of executive function, functional status, depression, and delirium using the Confusion Assessment Method (CAM).^25^ Executive function was assessed through two well-validated and widely used measures, Trail Making Test A (TMTA) and Trail Making Test B (TMTB).^26^ These tests require participants to connect a series of circles in ascending order. TMTA is composed of 25 encircled numbers; TMTB alternates between 23 encircled numbers and letters. Scoring is based on time to completion; a longer completion time indicates worse executive function. Raw scores were used in the analysis. Functional ability was assessed using the Instrumental Activities of Daily Living (iADL).^27^ Depression was assessed with the Geriatric Depression Scale (GDS), which is composed of 15 yes/no questions and is widely validated in the older adult population.^28^ Demographic information included age, sex, years of education, and tobacco use history. Data extracted from the electronic health record included preoperative blood pressure values, height, weight, past medical history including comorbidities, current outpatient medication use, and the American Society of Anesthesiologists Physical Status (ASA) classification for the planned surgical procedure. ASA classification ranges from 1-6 and is determined by the attending anesthesiologists prior to surgical intervention.^29, 30^ Each level has a specific definition with higher levels indicating increasing levels of disease, comorbidities and risk. As vascular surgery is often associated with delirium,^31^ it was selected *a priori* to be included as a covariate to examine whether surgical type was sufficient to predict delirium or if the additional information provided by a surgical risk score was necessary for a prediction model. Preoperative comorbidities as predisposing risk factors were assessed through the ASA score and Framingham Cardiovascular Disease 10-year Risk Calculator (Framingham CVD).^32, 33^ Framingham CVD was calculated through the online, interactive risk score calculator using data extracted from electronic health records and participant interview. This calculator includes sex, age, systolic blood pressure (SBP), treatment for hypertension, current smoker, diabetes and BMI. The most recent SBP value prior to surgery was used. BMI was calculated from collected height/weight values. The ACS NSQIP online surgical risk calculator^34^, (http://riskcalculator.facs.org) was used to obtain the risk scores for serious complications (NSQIP-SC) and death (NSQIP-D). Risk is calculated from both predisposing factors and precipitating factors (Figure 1). This calculator employs twenty patient predictors and pairs these with the Current Procedural Terminology (CPT) code, providing a risk score specific to each procedure. The inputted variables are as follows: age, sex, functional status (independent, partially dependent, dependent), emergency case, ASA classification, steroid use for chronic condition, ascites within 30-day prior to surgery, systemic sepsis within 48-hours prior to surgery, ventilator dependency, disseminated cancer, diabetes, hypertension with medications, congestive heart failure (within 30-days prior to surgery), dyspnea, current smoker (within 1-year), history of severe COPD, dialysis, acute renal failure, height and weight as well as surgical procedure. There are 1,557 distinct CPT codes, ranging from minor surgeries such as a cholecystectomy to major surgeries such as thoracoabdominal aortic aneurysm repair.

### Delirium Assessment

Pre- and postoperatively, participants were formally assessed for delirium and symptoms using the widely validated Confusion Assessment Method (CAM),^25^ 3D-CAM,^35^ and Delirium Rating Scale-R-98 (DRS),^36^ twice daily, between the hours of 0500-1000 and 1600-2200 on postoperative days 1-4 (regardless of day of the week). The CAM and 3D-CAM were administered concurrently to provide both a global and comprehensive view of delirium symptoms while providing a structured interview format. If the participant was CAM positive at the postoperative day 4 afternoon assessment, the participant was followed until delirium resolved to collect data on delirium duration. If participants were ventilated in the intensive care unit (ICU), the CAM-ICU^37, 38^ was administered. The DRS-R-98 (DRS) is a 16-item assessment tool that measures delirium symptoms and severity. The maximum score is 44-points, an increasing score indicates worse delirium.^36^ The research team (RDS, HL, ST, JK, LT, PR, MD, BH) met at least weekly to discuss delirium assessment findings and DRS ratings.

### Research Team

Each research team member (RDS, HL, ST, JK, LT, PR, MD, BH) underwent intensive training on delirium interview completion including CAM, 3D-CAM, CAM-ICU, and DRS. CAM training was completed as part of the NeuroVISION cohort study^39^ and the first author (HL) was officially trained on the CAM at the 2016 CEDARTREE Delirium Bootcamp and the CAM-ICU at the 2016 American Delirium Society pre-conference. Team members viewed 6 videos on CAM and 3D-CAM administration from the Hospital Elder Life Program^40^ website followed by an interactive team discussion on CAM/3D-CAM completion and observations. Team members shadowed the first author for six in-person CAM/3D-CAM assessments followed by in-depth discussion on observations, symptom rating (DRS), and bedside manner. The first author shadowed the team members for six CAM/3D-CAM/CAM-ICU/DRS assessments to ensure competency.

### Sample Size

The sample size was determined based on the need for a parsimonious delirium prediction model. We estimated three to four risk factors would form a reasonable clinical model if an AUROC >0.80 was obtained. Sample size was based on logistic regression and determined using the rule of 8-10 outcome events (delirium) per variable.^41, 42^ The decision to analyze was made after 100 participants were recruited with a delirium incidence rate of 32%.

### Statistical Analysis

Patient characteristics were described using means ± standard deviations for continuous variables and frequency counts with percentages for categorical variables. Dependent on the distribution of the data, continuous variables were compared using Student's t-test or Mann-Whitney U-test. Categorical variables were compared using *X^2^*. The outcome variables were Delirium Yes/No (DELYN) for logistic regression and delirium severity using the Peak DRS Total Score (DRS) for linear regression. Missing data was identified in the following variables (#missing): TMTB (1), GDS15 (1), TMTA (6), and Tobacco Pack Years (10). Little’s test of missing completely at random (MCAR) was not significant indicating that the missing data were missing completely at random and did not influence the analysis. Therefore, as outlined by Little (1988) the listwise deletion of participants with missing data was appropriate.^43^ Significance was notated with a p-value ≤0.05. NCSS v12.0, Stata/IC v15.0 and R v1.1453 were used for statistical analysis. These statistical packages were used to verify stable results across statistical software packages. HL, WC, and RB conducted the statistical analysis.

First, to evaluate the predictive ability of NSQIP-SC over NSQIP-D (as composites of the predisposing and precipitating factors), logistic (DELYN) regression models were completed and compared also to ASA classification and Framingham risk (as measures of predisposing factors only). Second, a delirium prediction model was developed. We did not employ univariate statistics to select candidate predictors as this may lead to poor performing predictors and overfitting.^44^ To counter the effects of small sample sizes and reduce bias within data, we employed a statistical shrinkage regression technique, using Least Absolute Shrinkage and Selection Operator (LASSO).^45-47^ This technique reduces the noise within the data, allowing true signals to be detected and avoids common problems such as model overfitting. Candidate variables demonstrating the smallest Mallow’s C*p* value,^48^ indicating precise predictors, were then applied in Best Subsets regression. Best Subsets regression is an automated regression approach that evaluates all possible combinations of candidate predictors.^49, 50^ The output provides a set models with model fit statistics. Model selection was based on assessment of model fit using Akaike information criteria (AIC), Bayesian information criteria (BIC), and McKelvey and Zavoina’s Pseudo-R^2^.^51^ The area under the receiver operating characteristic curve (AUROC) with 95% CI was calculated. Calibration was assessed through goodness-of-fit tests calculated by the Hosmer-Lemeshow statistic. Sensitivity, specificity, positive predictive and negative predictive values were calculated and reported.

The peak delirium severity (DRS) score was transformed using the Box-Cox Method^52^ with the optimal Lambda value due to the positive skew, please refer to Figure 4-A and B **for raw and transformed plots**. The regression modeling procedures outlined in the paragraph above for logistic regressions were repeated for the linear regression model. Model selection was based on assessment of model fit using Akaike information criteria (AIC), Bayesian information criteria (BIC), and adjusted R^2^.^51^

## Results

Thirty-one participants (32%) experienced postoperative delirium with a mean peak DRS severity total score of 21.48 (±SD 6.40). Forty-two percent experienced hypoactive delirium. The median delirium duration was one day (24 hours). Participant characteristics are summarized in Table 1. Delirious patients had higher preoperative NSQIP risk scores of serious complications (NSQIP-SC) and death (NSQIP-D), worse executive function tests, were more likely to have had a vascular surgery, and higher ASA status (univariate, p<0.05). Significant pairwise correlations were demonstrated between the DRS and NSQIP-SC, NSQIP-D, TMTA and TMTB (univariate, p<0.05). No significant differences were identified between the non-delirious and delirious groups in terms of age, sex, education level, past/present tobacco use, blood pressure metrics, functional status, GDS15, and Framingham CVD.

**Table 1:**
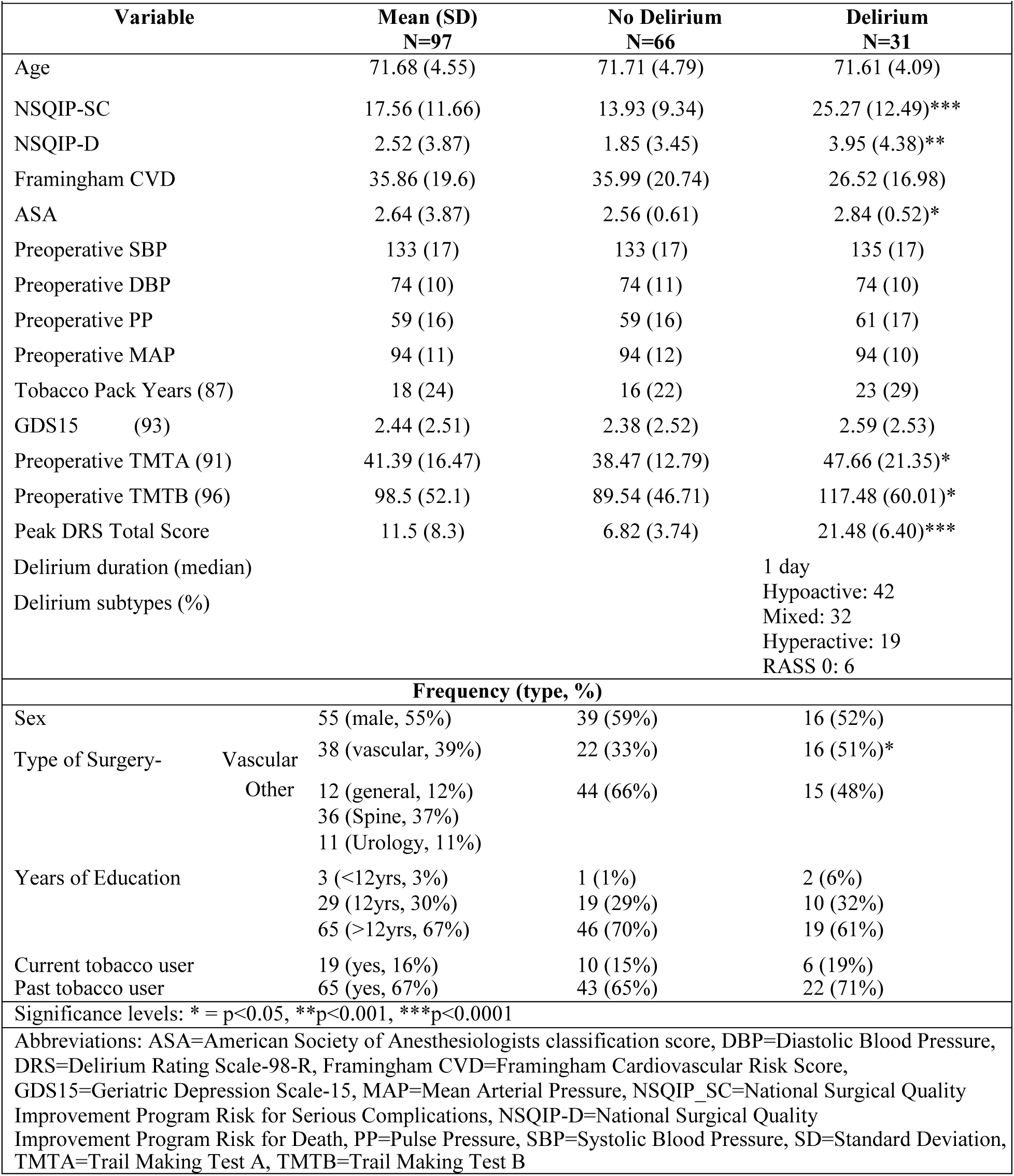
Description of sample and significant differences between no delirium and delirium

### Surgical Risk Scores and Delirium Incidence

We confirmed preoperative NSQIP-SC as a robust predictor of postoperative delirium incidence using single factor logistic regression models for NSQIP-SC and NSQIP-D. Both demonstrated moderate to fair predictive ability with an AUROC of 0.76 (95% CI: 0.66, 0.87) and 0.73 (95% CI: 0.62-0.84), respectively. Although similar in their projected ability to predict individuals at a higher risk for delirium development, support for the NSQIP-SC model, over the NSQIP-D model, is provided by optimal AIC (1.086) and the BIC (−333.254) metrics. NSQIP-SC also performed better than the ASA classification (AUROC 0.63, 95% CI: 0.53-0.73) and the Framingham risk score (0.53, 95% CI: 0.40-0.65). The reported odds ratios, coefficients, AUROC, sensitivity and specificity, and model fit statistics including AIC and BIC are in **Supplementary Table 1**.

### Derivation of a Delirium Incidence Prediction Model

In order to derive a potential delirium incidence prediction model we followed TRIPOD guidelines to apply a statistical shrinkage technique (LASSO) followed by Best Subsets regression using age, sex, NSQIP-SC, NSQIP-D, tobacco pack years, vascular surgery, ASA classification, Framingham CVD, GDS15, TMTA, TMTB, and blood pressure metrics. A two-factor logistic regression model containing preoperative NSQIP-SC and TMTB to predict postoperative delirium incidence was selected as this model demonstrated optimal model fit statistics. Hosmer-Lemeshow goodness-of-fit test was not significant (p=0.37), indicating accurate model calibration. The model demonstrated moderate predictive ability (AUROC 0.81, 95% CI: 0.72-0.90), a 5-point increase in the NSIQP-SC score increased the probability of delirium incidence by 10% (Figure 3). The logistic regression model including classification metrics is displayed in Table 2.

**Figure 3.**
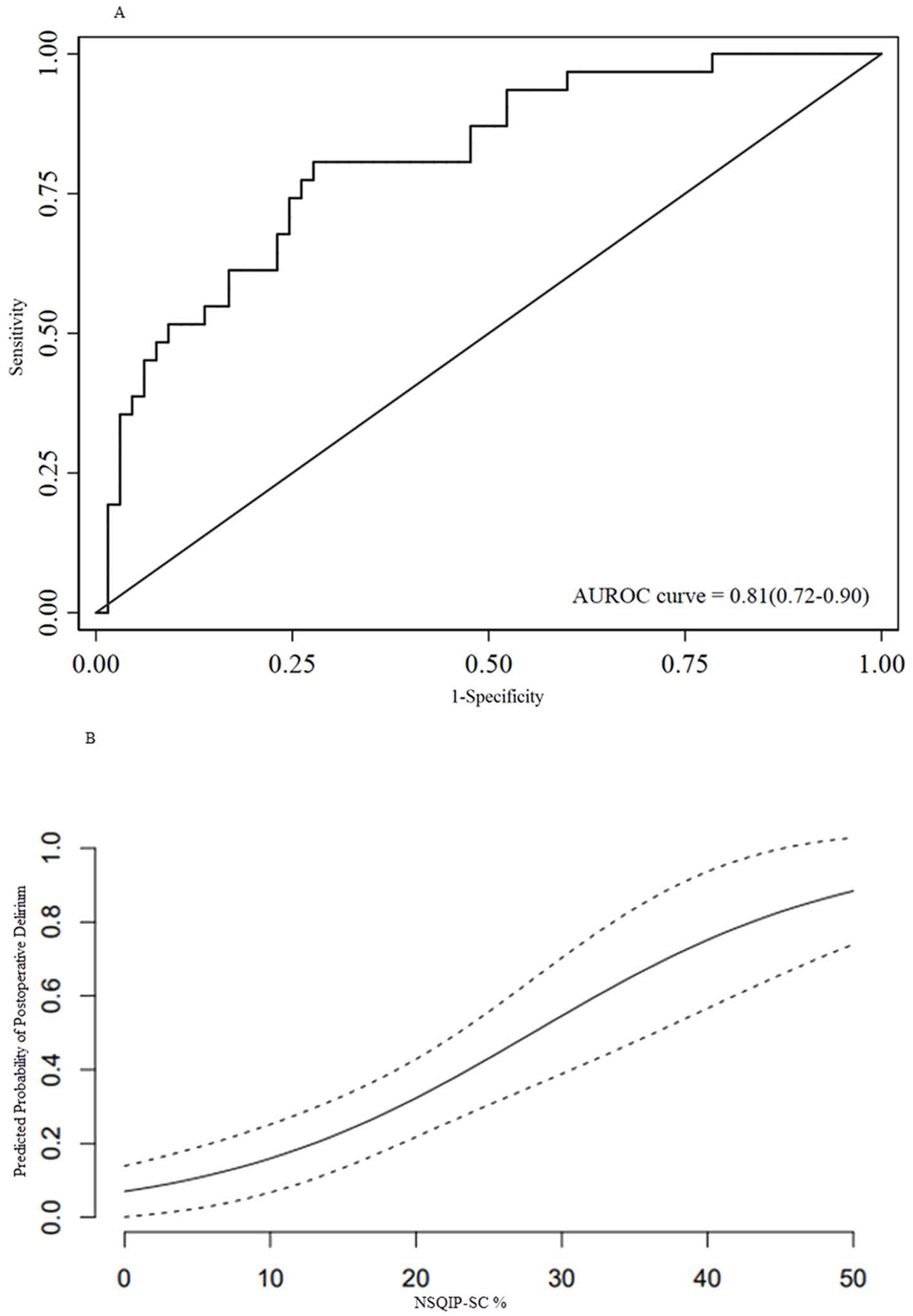
Illustrates the predictive ability of the NSQIP-SC and TMTB model for postoperative delirium incidence. (A) Displays the Area Under the Receiver Operator Curve statistic (AUROC). (B) Demonstrates the predicted probability of postoperative delirium incidence based on the % NSQIP-SC score. This is holding TMTB constant at zero.

**Table 2:**
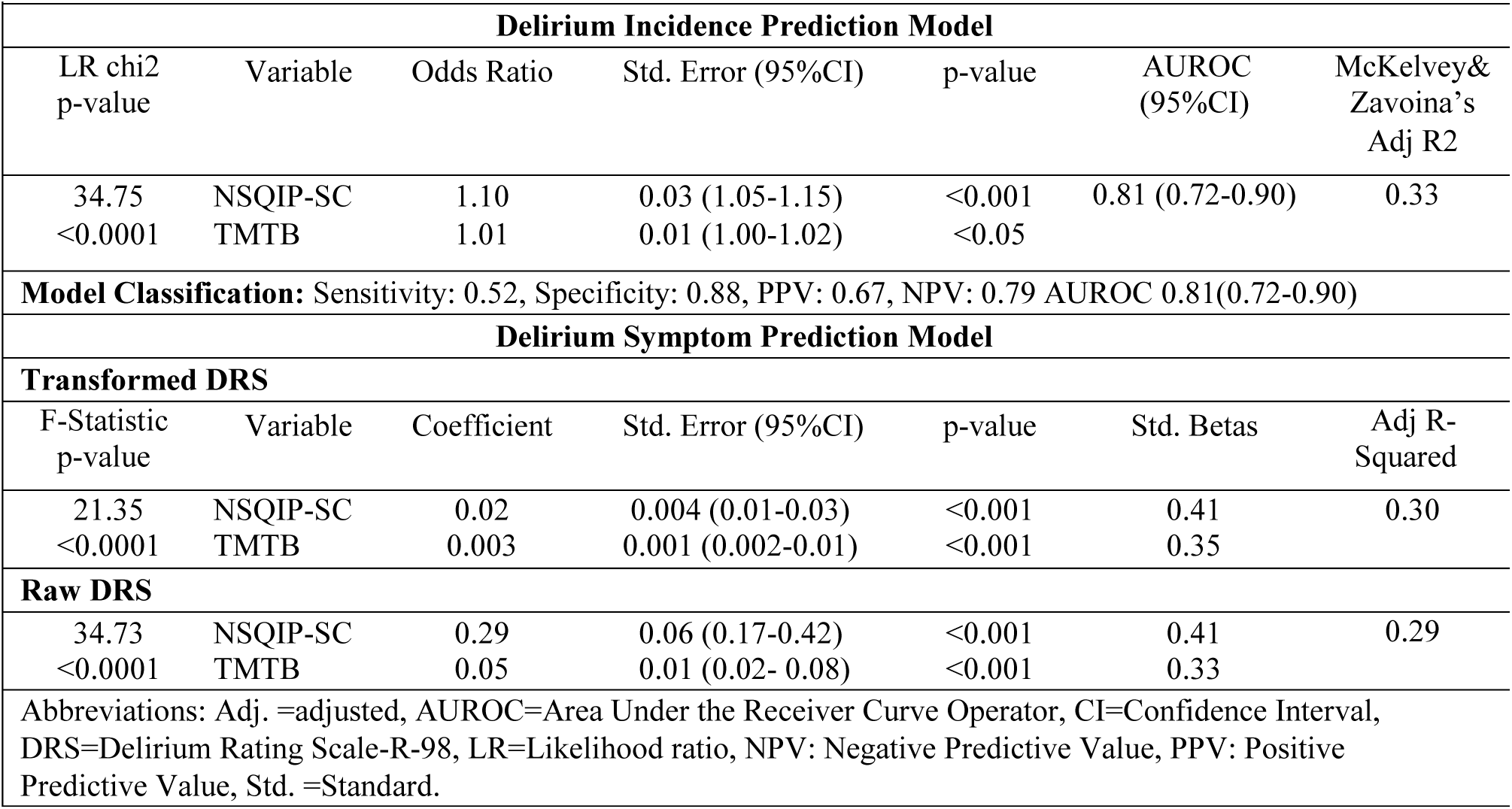
Results of LASSO and Best Subsets Regression. Model Statistics shown below.

### Derivation of a Delirium Symptom Prediction Model

Preoperative NSQIP-SC was confirmed as a predictor of postoperative peak DRS using simple linear regression models for NSQIP-SC (p<0.0001, AdjR^2^: 0.184). Similar to the logistic regression results above NSQIP-D was also significantly associated with DRS (p=0.04, AdjR^2^: 0.03). However, the NSQIP-SC model demonstrated higher adjusted R^2^, and lower AIC (1.525) and BIC (−290.718) metrics, providing support for that predictor over NSQIP-D. Similar to logistic regression for postoperative delirium incidence, ASA (p=0.04, AdjR^2^: 0.03) and Framingham risk score (p=0.69, AdjR^2^: 0.01) did not demonstrate a strong predictive relationship with DRS.

The model demonstrating optimal fit statistics for postoperative peak delirium severity was a two-factor linear regression model containing preoperative NSQIP-SC and TMTB using LASSO and Best Subsets regression. This two-factor model reports an adjusted R^2^ of 0.30 (p<0.001), thus explaining 30% of the variability in observed delirium symptoms. For every 1-standard deviation (SD) increase in the NSQIP-SC score, the DRS SD will increase by 0.42 points. Further model details are displayed in **Table 2** and Figure 4. Age, sex, NSQIP-D, ASA, tobacco pack years, vascular surgery, Framingham CVD, GDS15, TMTA, and blood pressure metrics were not identified as significant predictors.

Figure 4

Illustrates the postoperative delirium symptom severity prediction model. Box A is a histogram showing the data distribution of the Peak Delirium Rating Scale Score (DRS). This value was transformed using the Box-Cox Method with an optimal lambda value of 0.35 achieving a near Gaussian distribution and is shown on the histogram in Box B. Boxes C-E display the predicted burden of delirium symptoms based on the NSQIP-SC and TMTB prediction model (Box C) and univariate analysis of NSQIP-SC (Box D) and TMTB (Box E). The statistics from each regression model are shown in the upper left hand corner of each box. The univariate NSQIP-SC regression model was analyzed with 97 participants. Due to one missing assessment of TMTB, Box C and E are analyzed with 96 participants.

**Figure 4-1.**
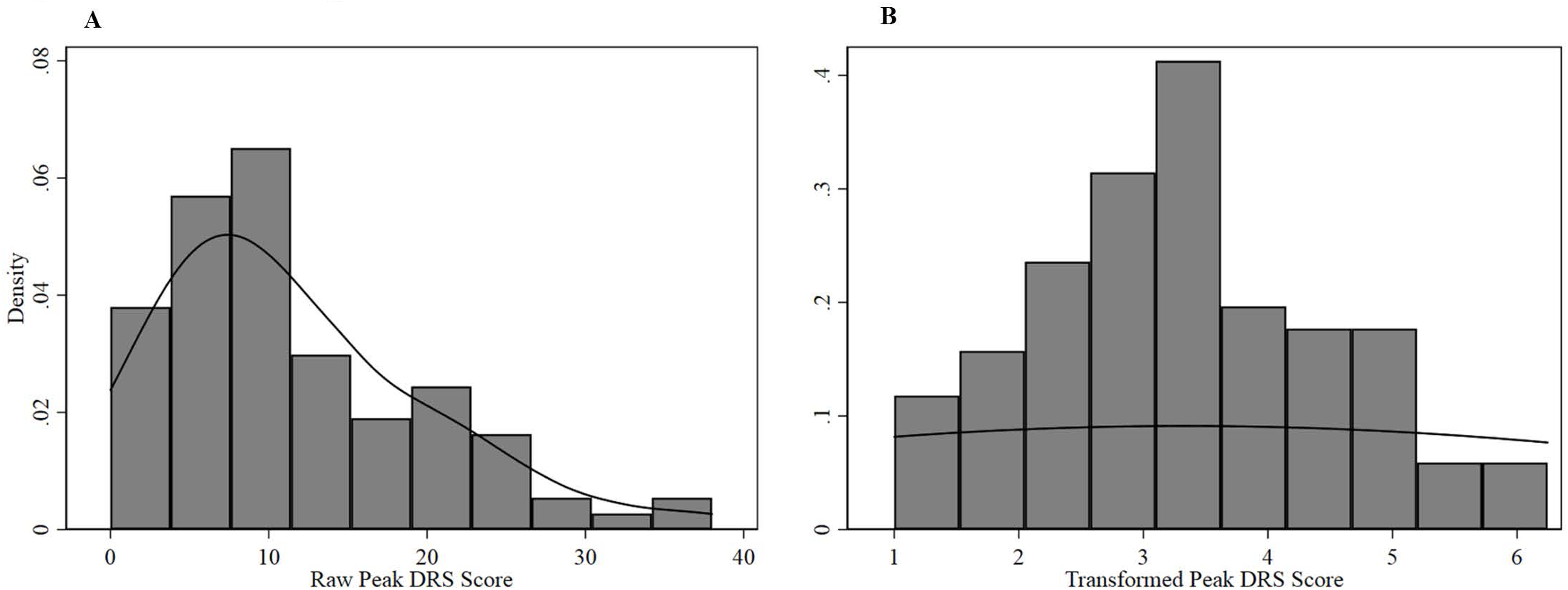
Peak DRS Histograms

**Figure 4-2.**
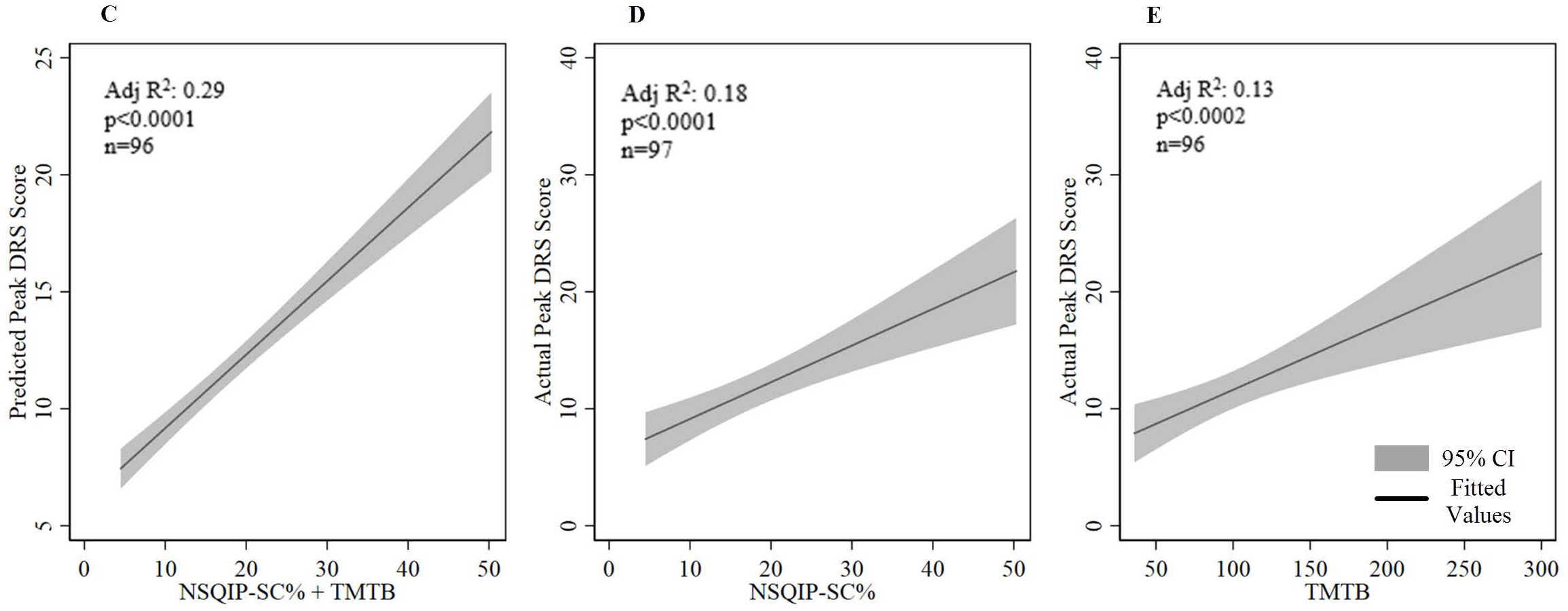
Delirium Severity Prediction Model

## Discussion

Our analysis of a prospective perioperative cohort found NSQIP-SC to be a robust predictor of postoperative delirium incidence and severity. The preoperative NSQIP-SC score combines data on both predisposing risk factors and the estimated magnitude of the surgery, i.e. the precipitating event and is well positioned to contribute information on the risk of delirium. The addition of the executive function measure, TMT-B, improved performance of both prediction models. Previous research has identified executive function to be significantly associated with delirium incidence.^22-24^ However, as identified by our recent systematic review of delirium prediction models in older adults, an executive function measure has not been applied previously.^12^ Given the breakdown in executive function during delirium and prior data on the predisposition to delirium by impaired cognition, incorporating a cognitive variable appears biologically important; our data show that it is statistically important too. This study expands current knowledge by examining the utility of NSQIP risk scores and executive function in predicting delirium incidence and severity.

The NSQIP-D score was significantly associated with both postoperative delirium incidence and severity, however, it was not selected as a predictor in our variable selection. Delirium often results from a complicated perioperative course, particularly following a major surgery, hence a relationship with NSQIP-SC is plausible. Furthermore, a recent systematic review questioned the strength of the association between postoperative delirium and mortality, hence *a priori* we hypothesized that NSQIP-SC would perform better than NSQIP-D for predicting postoperative delirium incidence and severity.^20^ This was supported by several statistical measures in our dataset. The modeling procedures, LASSO and Best Subsets regression, did not identify vascular burden, comorbidities, smoking history, and depression as important predictors although these variables have been identified as significant risk factors for postoperative delirium incidence in prior research. This may be due to a number of factors. First, their prevalence in this population of study may not be sufficient for prediction. In order for a risk factor to also be an accurate predictor, it must be sufficiently prevalent in the at-risk population.^53, 54^ Secondly, late-life depression, vascular burden, and tobacco use often co-occur leading to overlapping data capture. Variables that capture similar information fail to contribute important information during modeling procedures.^12^ Lastly, these variables contribute information on predisposing risk, but do not include valuable information on the future precipitating event, the planned surgical procedure. The inclusion of information about the future precipitating event, as done in NSQIP-SC, contributes crucial information to the prediction of postoperative delirium.

### Strengths

The strengths of this study include its prospective perioperative design, statistical methods chosen, and rigorous delirium assessments including outcomes based on incidence and severity of delirium. This novel application of a readily available, simple tool has potential for broad application in delirium-focused clinical care. The NSQIP-SC score combines several potential preoperative risk factors for delirium (age, functional status, current tobacco use, vascular burden) with the precipitating event, the planned surgery, and provides a single risk score that is easy to interpret, *i.e.*, a 5 point increase in NSQIP-SC results in a 10% increase in the probability of a patient experiencing delirium. Given that the patient becomes delirious postoperatively, quantifying the potential impact of this precipitating event is clearly a key feature of a delirium prediction model.

Delirium represents a severe breakdown in an individual’s executive function. The *Cognitive Disintegration Model* posits that there is a critical threshold (i.e. the “Delirium Threshold”) in cognition (or network connectivity) that must be crossed for delirium to result. Individuals with more vulnerability (predisposing factors) going into surgery, i.e. older age, multiple comorbidities and worsening executive function, are closer to that “Delirium Threshold” (Figure 1) and may need a smaller precipitating event to push them into delirium. In this context, inclusion of executive function, which is not captured in NSQIP-SC, improved model performance. Furthermore, this simple two-factor model captures information on both the predisposing and precipitating factors for delirium. Its parsimonious nature is a clinical strength.

### Limitations

This study has several limitations to consider. While these models were built using statistical methods optimized for modeling a limited number of events, the small sample size may still have had an effect on the results. The recommendations by the TRIPOD guidelines and the CHARMS checklist were followed to use statistical shrinkage procedures to minimize model overfitting. Nonetheless, it is worth emphasizing the study sample is small with only 31 delirious participants. The population is largely homogenous in terms of years of education and ethnicity. In larger and more diverse populations, additional factors may enhance model performance. Future studies should focus on the broad external validation of these models following the statistical methodology outlines by the TRIPOD guidelines. This requires a new perioperative cohort study that will recruit patients from diverse populations and will be the target of future grant applications.

### Conclusion

This analysis of a prospective perioperative cohort study identified NSQIP-SC and TMT-B as predictors for delirium incidence and severity. A preliminary delirium prediction model was created for both delirium incidence and severity. These models should be validated in future studies.

## Details of Authors Contributions

Patient recruitment: HL, ST, PR, JK, MD, LT, BH, RDS

Clinical execution: HL, ST, PR, JK, MD, LT, BH, RDS

Guidance on cognitive tests: LB

Data analysis and interpretation: HL, WC, RB, RDS

Manuscript writing: HL, RDS

Manuscript revision: LB, ST, PR, JK, MD, LT, BH, WC, RB

Study Conception, protocol: RDS

Study idea: HL, RDS

## Acknowledgements

Much gratitude to those that provided guidance and support throughout the conduct of this study.

Collected data: Mitch Whalen, Michelle Prihoda, Maggie Schmit, Casandra Stanfield, and Daniel Wayer.

Supported study: Staff and clinicians at University of Wisconsin Hospital and Clinics Critically reviewed the proposal: Drs. Tonya Roberts, Kirk Hogan, Kris Kwekkeboom. David Dwyer

